# Mathematical modeling of clonal interference by density-dependent selection in heterogeneous cancer cell lines

**DOI:** 10.1101/2023.05.08.539618

**Authors:** Thomas Veith, Saeed Alahmari, Andrew Schultz, Joseph Johnson, Konstantin Maksin, Noemi Andor

## Abstract

Many cancer cell lines are aneuploid and heterogeneous, with multiple karyotypes co-existing within the same cell line. Karyotype heterogeneity has been shown to manifest phenotypically, affecting how cells respond to drugs or to minor differences in culture media. Knowing how to interpret karyotype heterogeneity phenotypically, would give insights into cellular phenotypes before they unfold temporally. Here we reanalyze single cell RNA (scRNA)- and scDNA sequencing data from eight stomach cancer cell lines by placing gene expression programs into a phenotypic context. We quantify differences in growth rate and contact inhibition between the eight cell lines using live-cell imaging, and use these differences to prioritize transcriptomic biomarkers of growth rate and carrying capacity. Using these biomarkers, we find significant differences in the predicted growth rate or carrying capacity between multiple karyotypes detected within the same cell line. We use these predictions to simulate how the clonal composition of a cell line will change depending on the timing of splitting cells. Once validated, these models can aid the design of experiments that steer evolution with density dependent selection.

## 1 INTRODUCTION

Cellular heterogeneity is a defining feature of most cancers and critical to tumor progression and treatment failure [1, 2]. Advances in sequencing techniques have provided for an unprecedented depth of genetic profiling and ushered in a new era of individualized, data-driven cancer genetics [3]. Despite this progress, the relationship between genetic and phenotypic heterogeneity remains a significant gap in our current understanding.

One contributor to a cancer cell’s phenotype are large-scale somatic copy number alterations (SCNAs) of 10 mega base pairs or more. Studies show that SCNAs correlate with progression free and overall survival, with cancer cells grouped by copy number landscape exhibiting the same resistance to chemotherapy [4, 5, 6, 7]. SCNAs of whole chromosomes or chromosome arms, also known as aneuploidy, is the most common characteristic of malignant cells [8]. Variation in copy number arises mainly from chromosomal instability (CIN), a hallmark of cancer [9]. CIN has been shown to promote metastasis and tumor evolution, particularly in cancers which have common aneuploidy patterns, such as gastric cancer [2, 10]. Studies have shown that SCNAs activate oncogenes and disrupt tumor suppressor genes [11], correlate with cancer phenotype [12, 13, 14, 15], and are spatially segregated within the tumor [16]. Aneuploidy fuels rapid phenotypic evolution and drug resistance, with colonies resistant to the same drug displaying similar karyotype profiles [17]. However, how selection acts upon SCNAs remains poorly understood.

Selective pressures naturally vary in the tumor over space and time. For example, a cell that is in a very densely packed environment will be under more pressure to overcome contact inhibition than a cell that finds itself in sparse conditions. This variation in evolutionary pressures likely contributes to the coexistence of cancer cells with heterogeneous phenotypes. Density dependent selection occurs when fitness is a function of population density [18]. A related concept in ecology is life history theory [19], and the r/K-selection framework, which investigates trade-offs between the number of offspring a species produces (growth rate or ‘r’) and the ability to compete in dense ecological niches (carrying capacity or ‘K’) [20, 21].

Here we present a framework for examining how density dependent selection acts on coexisting clones defined by SCNAs and apply it to a set of stomach cancer cell lines. Hereby we focus on carrying capacity as defined by spatial limitations, rather than metabolic constraints.

## 2 RESULTS

### 2.1 Identifying biomarkers of growth rate and contact inhibition

Sequencing the DNA and RNA of >36,000 cells from nine stomach cancer cell lines, our prior work classified cells into groups with unique karyotypes [22], further referred to as superclones [23] or clones. To compare growth dynamics across cell lines we grew eight of these stomach cancer cell lines in a T25 flask until they reached confluence (6-23 days), imaging them every day to count cells (Online Methods). Comparing Countess derived cell counts to cell counts derived from live cell imaging confirmed segmentation accuracy (Supplementary Fig. 3), supporting feasibility of monitoring growth dynamics during routine *in-vitro* experiments.

We fit the Gompertz growth model and two instances of the generalized logistic growth model (Richards and Verhulst) [24] to this time-series data (Fig. 1A; see also Online Methods 4.5). The resulting *R*^2^ was high (Adj-*R*^2^ > 0.95) for each of the models and cell lines (Supplementary Table 1). Using Akaike Information Criterion, we determined that Richards model slightly outperforms Verhulst, which in turn slightly outperforms Gompertz (Supplementary Table 5). We thus eliminated Gompertz from consideration for model selection. Using likelihood profiling [25] to assess practical identifiability, we concluded that noise levels in data collected for 4/8 cell lines are too high to infer all parameters of the Richards model (Supplementary Table 6). Therefore, we used the simpler Verhulst model (commonly referred to as the “logistic function”) to infer growth dynamics from time-series data for all eight cell lines (see Fig. 1, Supplementary Table 2).

**Figure 1.**
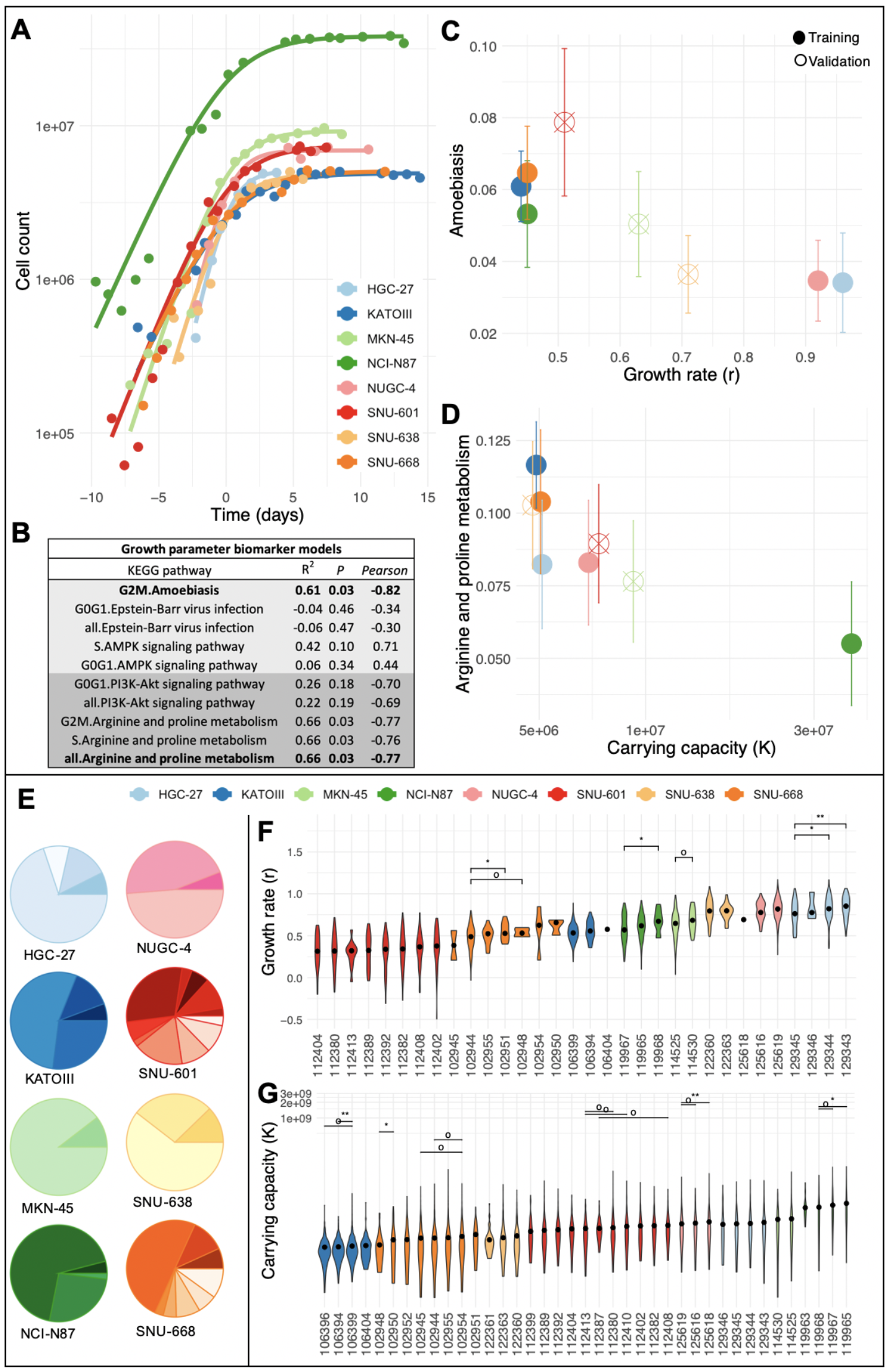
Identifying biomarkers of growth and carrying capacity. **(A)** Logistic growth curves fit to the cell counts of eight gastric cancer cell lines at various stages of their growth. Fits are shifted along the x-axis such that the midpoint of each curve lies above *x* = 0. The eight cell lines differ in their maximum growth rate (*r*) as well as their maximum sustainable population size (*K*). (**B**) Summary statistics of linear regression models used to correlate KEGG pathway activity levels with logistic function growth parameters. The top five models and their performance in the training cell lines are shown for growth rate (*r*: light gray rows) and carrying capacity (*K*: dark gray). Columns: Pearson = Pearson correlation coefficient; P = FDR corrected p-value; *R*^2^ = adjusted *R*^2^. (**C-D**) Relation between growth parameters (*r, K*) and the model with best performance in the validation data set: ‘Amoebiasis’ **(C)** and ’Arginine and proline metabolism’ **(D)**, respectively. Pathway activity quantified using AUCell with scRNA-seq data. Error bars represent median absolute deviation. **(E)** Clonal composition confirmed by both scDNA- and scRNA-seq in eight gastric cancer cell lines (data taken from [22]). (**F-G**) Violin plots showing predicted values for clonal growth rate as a function of ’Amoebiasis’ activity (**F**), and clonal carrying capacity as a function of ’Arginine and proline metabolism’ activity (**G**). Wilcoxon signed-rank test: o P<0.1, * P<0.05, ** P<0.01, *** P<0.001. If a cell line has more than three clone pairs with significant differences, we display only the three highest p-values below 0.1.

Overall, there was no significant correlation between growth rate and carrying capacity across cell lines (Pearson’s *r* = -0.37, *p* = 0.54). In order to investigate potential r/K trade-offs between clones within a cell line, we sought to find transcriptomic biomarkers of the inferred growth parameters in scRNA-seq data available from the prior study [22]. Recently, investigators induced r/K-selection in HeLa cells [26, 20]. The authors found that genes which were differentially expressed between r and K-selected cells cultured at low densities, were enriched among 25 pathways defined in the KEGG database. We tested these pathways for their potential as biomarkers of growth rate and carrying capacity in five of the eight gastric cancer cell lines (further referred to as training set; Supplementary Table 3). For each of the 25 pathways, we fitted a linear regression model to predict the growth parameters inferred for a given cell from the cell’s respective pathway activity level. Because the cell cycle is one of the strongest modulators of pathway activity [22, 27, 28], we also grouped the cells according to their assigned cell cycle state, calculating the median growth parameter and pathway activity across: (i) G0G1 cells, (ii) S cells, (iii) G2M cells and (iv) all cell cycle states combined. The pathways which had the strongest predictive value for growth rate (*r*) included *’Amoebiasis during G2M’* (adj-*R*^2^ = 0.89,*P* < 0.02), *’Epstein-Barr virus infection’*, and *’AMPK signaling pathway during S-phase’*. For carrying capacity (*K*), *’PI3K-Akt signaling pathway’* (adj-*R*^2^ = 0.91,*P* < 0.01) and *’Arginine and proline metabolism’* had the strongest signal (Supplementary Table 3). This process was repeated, now including the remaining three cell lines (further referred to as validation set), but only for these top performing pathways and cell cycle states (Fig. 1B). Of these top pathways, 40% were confirmed in the validation set (FDR adjusted *P* < 0.05; Fig. 1B), including *’Amoebiasis during G2M’* and *’Arginine and proline metabolism’*, which were further used as proxies of growth rate (*r*) and carrying capacity (*K*) respectively.

### 2.2 Towards informing future experiments: steering clonal evolution

Identification and characterization of clones within a cell line requires high-throughput assays, such as single-cell sequencing, rendering repeated measurements of clonal composition at a high temporal resolution cost-prohibitive. This underscores the need to identify biomarkers that can be used to estimate growth parameters.

We previously identified 39 clones, each defined by distinct SCNAs and confirmed by both scDNA- and scRNA-seq, across the eight cell lines [22] (Fig. 1E). Using pathway activity levels from the KEGG pathways *Amoebiasis* and in *Arginine and proline metaboism*, we predict growth rate and carrying capacity for every sequenced cell in each cell line. Grouping cells by their clone membership (Fig. 1F-G) we conclude that up to 2 r/K trade-offs may exist between clone pairs in two of the eight analyzed cell lines (Fig. 2A; Online Methods and Supplementary Table 4).

**Figure 2.**
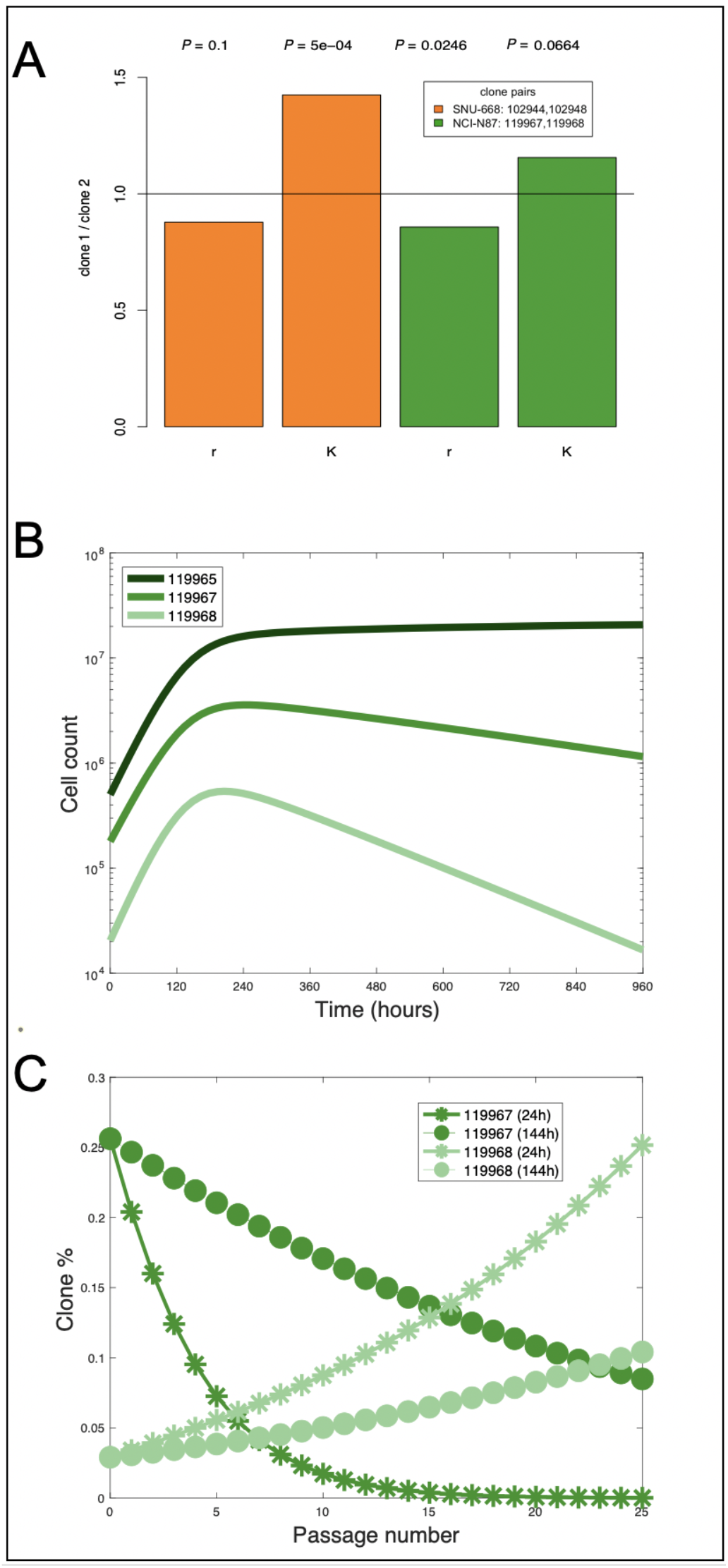
Using biomarkers of growth and contact inhibition to steer clonal evolution across passages. **(A)** Clone pairs with strongest difference in both growth rates (*r*) and carrying capacities (*K*) are detected in two cell lines. Each color encodes for a single pair of clones (legend). Each bar is calculated as the ratio of *r* or *K* between the pair of clones shown in the legend. Horizontal line at 1 indicates identical parameters for both clones. Clone pairs with potential r/K trade-offs will be represented by bars on both sides of the horizontal line. P-value of difference in *r* or *K* between a given pair of clones is indicated on top of each bar. **(B)** Simulated growth of the three largest clones (color-code) detected in the NCI-N87 cell line over 40 days (smallest clone excluded due to insufficient G2M cell representation). **(C)** Change in frequency of the two clones shown in (**A**) over multiple passages (x-axis). By changing when to split the cells after each passage (stars vs. circles in legend), we can accelerate or delay the decline of clone 119967 (dark color shades), from passage 6 (intersection of star-shaped curves) to passage 23 (intersection of circle-shaped curves). A seeding density of 700,000 cells was used as initial condition. Clonal frequencies at harvest set the initial frequency conditions for each subsequent seeding.

One of these potential r/K trade-offs was in the NCI-N87 cell line. Optimizing the timing for splitting cells should thus enable evolutionary steering of the cell line’s clonal composition across passages. A condition for such an optimal time to exists is that growth of the population can never be negative, which we prove analytically (Supplemental Section 2). We used growth parameters predicted for the clones identified in the NCI-N87 cell line (Supplementary Table 4) to parameterize a multi-compartment ODE model to simulate how clonal composition changes over time (Fig. 2B):

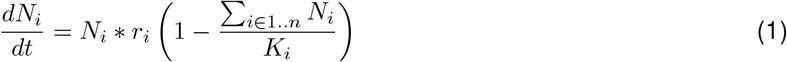

where *N*_*i*_ is the cell representation of clone *i. K*_*i*_, *r*_*i*_ are the biomarker-inferred carrying capacity and maximum growth rate of each clone respectively (see also Online Methods 4.7).

Simulations shown in Fig. 2B predict that shifts in clonal composition will likely occur as NCI-N87 cells grow *in-vitro*, with the direction and magnitude of these shifts depending on the timing of splitting cells (Fig. 2C). For example, changing when to split the cells after each passage, we predict can accelerate or delay outcompetition of NCI-N87’s clone 119967 by clone 119968 for 17 passages (Fig. 2C). This suggests that for a subset of cell lines the timing for splitting cells can be optimized to either accelerate or delay changes in clonal composition.

## 3 DISCUSSION

Through a reanalysis of existing scRNA-seq data from eight stomach cancer cell lines [22], we confirm biomarkers of growth rate and contact inhibition previously identified in breast cancer cell lines [20], suggesting similarities in the mechanisms of contact inhibition exist across multiple tissue and cancer types. The strongest biomarker of growth rate (’Amoebiasis’) is commonly upregulated in gastric cancer patients with poor outcomes [29, 30, 31]. ’Arginine and proline metabolism’ – the strongest biomarker of carrying capacity – reflects metabolites known to be abnormal in gastric cancer patients [32]. Arginine blood levels have been shown to have both diagnostic and prognostic value [32, 33]. Moreover, arginine blood levels are higher in primary tumors compared to metastatic lesions, and high-arginine patients have better overall survival than low-arginine [32]. Proline metabolism has been shown to differentiate between gastric cancer subtypes with proline metabolism being upregulated in the non-CIN subtype [34]. A separate study has shown that one of our cell lines (HGC-27) upregulates ’Arginine and proline metabolism’ when subjected to simulated microgravity, which has potentially interesting implications for the effects of contact inhibition on metabolite preference [35]. Regulation of this pathway also plays a role in activating AMPK/mTOR pathways to promote progression in gastric cancer [36].

Differential expression of biomarkers for growth rate and contact inhibition/carrying capacity between clones within a cell line (Fig 1), suggest that different population densities will alter a cell line’s clonal composition. Indeed, we previously observed changes in clonal composition over only 5-7 passages in a subset of these cell lines [22]. Another recent study profiled genome-wide copy numbers and mRNA levels in HeLa cells and observed progressive divergence over 50 successive passages [37]. Recent reports examined the responsiveness of 21 different variants of the same cell line with 321 anti-cancer agents. They found that 75% of compounds which strongly inhibited some variants were inactive in others, and their results suggest phenotypic heterogeneity is associated with genetic variation due to clonal evolution [38]. These studies emphasize that every cell division is an opportunity for the cells to mutate and adapt to their environment and underscores the need for routine tracking of the pedigree of evolving cell populations over decades along with potentially changing environments (e.g. cell culture habits, therapy). Such efforts could help reveal long-term trends in the evolution of cell lines, that remain elusive at shorter time-scales.

## 4 ONLINE METHODS

### 4.1 Cell culture

The identity of each cell line was determined through independent karyotyping and mycoplasma contamination assessed. Cells were cultured in their recommended media conditions at 37C. For, HGC-27, EMEM (Quality Biological Inc); KATOIII, RPMI-1640 and EMEM (1:1 mix); NCI-N87, RPMI-1640 (ATCC modified); the remaining five cell lines (MKN-45, NUGC-4, SNU-601, SNU-638, SNU-668) used RPMI-1640. All cell lines were grown in the aforementioned media with 10% fetal bovine serum (Gibco) and 1% penicillin–streptomycin (Gibco). For cell passaging, Trypsin EDTA (0.25%) with phenol red (Gibco) was added, followed by inactivation using the respective growth media.

### 4.2 Microscopy

Cells were seeded in a T25 flask (Fisherbrand), and sets of 4 phase contrast images were taken at 20X magnification for NCI-N87, and 10X magnification for the aforementioned cell lines on an Evos FL. Segmentation using Cellpose was utilized to quantify the cell count and cellular features (number of detections, centroid X *μ* m, centroid Y *μ*, ROI, area *m*^2^, perimeter *μ*m). Growth curves were generated to examine the carrying capability of each cell line.

#### Image preprocessing

The microscopy image acquisition and light settings could results in variations on the brightness of acquired images. The automatic segmentation and counting pipeline applied to dark images showed inaccurate results (i.e, mostly false negatives). Therefore we applied pre-processing steps to our pipeline, consisting of: (i) gamma correction to reduce darkness; (ii) histogram equalization to improve image contrast and (iii) Gaussian blurring for smoothing.

#### Cell segmentation

Fully automated cell segmentation of phase contrast images was performed using Cell-ose software. Cellpose is a deep learning approach based on the U-Net architecture for cell segmentation [39][40], where vertical and horizontal spatial gradients of cells are predicted. Furthermore, Cellpose predicts a binary map of a cell location either inside or outside a region of interest (ROI). Using both the combined vertical and horizontal gradients and the binary map, cell localization and generation of binary masks for every cell are done. Our pipeline for cell segmentation using Cellpose uses 2D microscopic images as input. An ROI annotation rectangle is applied on the phase contrast image (left corner index of the rectangle: (100,100); rectangle width and height: (1800px, 1100px). This ROI is used for subsequent image segmentation and analysis. We fine-tuned a pre-trained Cellpose model called (Cytotorch_2) for learning to segment cells on given microscopic images of different cell-lines. After learning the segmentation using Cellpose, the trained model was tested on a hold-out set for evaluation. Then, feature extraction of each segmented cell was performed for further analysis of cell growth at a given time point. The features extracted from each cell segmentation include area, perimeter, roundness, and centroid. The pipeline saves the extracted features and visualization of the segmentation onto a user-desired folder for cell growth estimation and analysis.

### 4.3 Determination of clonal growth parameters

Determining relative clonal carrying capacities, growth rates, and loss of contact inhibition parameter values is a necessary first step in modeling density dependent selection in heterogeneous cell lines. To achieve this we use gene expression signatures as surrogate of growth parameters as previously described [41, 42].

#### Quantification of single cell pathway activity from gene expression

In order to identify cells with active gene sets, we utilized AUCell version 3.18 for R version 4.3.1 [43]. AUCell takes as its input the scRNA-sequencing data for the cells of interest and a list of gene sets. The output is the gene set activity for each cell. AUCell uses the area under the curve across the rankings of all genes of a particular cell, where genes are ranked by their expression value. A rank-based scoring method means AUCell is not affected by units or normalization method of the gene expression data. Genes in the top 5% of the ranking are considered active in a given gene set. In order to account for potential batch effects, scRNA-sequencing data for all cell lines was analyzed together. Cells with less than 200 features were excluded. Seurat version

2.3.4 was used to create a Seurat object for input to AUCell to quantify the activity of more than 2000 pathways from the KEGG database. For all further analysis, we focused on only a subset of 25 KEGG pathways, which were previously identified as being differentially expressed between r-selected and K-selected HeLa cells [20].

#### Clonal growth parameters

We tested these 25 KEGG pathways quantified by AUcell as biomarkers of growth rate (*r*) and carrying capacity (*K*). The test takes the form of a linear model:

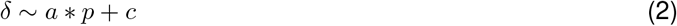

where *δ* is the parameter value (*r* or *K*) and *p* is the pathway activity. The pathways are then ranked by adjusted-*R*^2^ and the top five best-fitting pathways are prioritized as potential biomarkers of a given parameter. With the linear models correlating pathway activity with growth parameters built at the cell line level, we can predict growth parameters for all clonal populations using their pathway activity as input (Fig. 1E-F, Supplementary Table 4). Once parameters have been predicted, clonal growth can be simulated as systems of paired ODEs (see section 4.4).

### 4.4 Mathematical models of *in-vitro* cell growth

The nature of tumor growth is not well-known, and the exact laws which govern the growth of tumor cells will likely be context-dependent (cancer type, location in the body, etc). However, even when many of these dependencies are held constant when cells are grown in a dish, it is difficult to discern between various models of cancer cell growth [44]. We fit generalized logistic growth models [24] to the time-series cell growth data for eight gastric cancer cell lines (Supplementary Table 1):

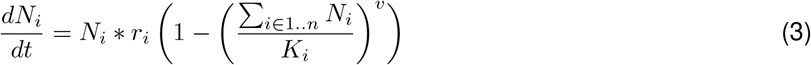

where *i* is either the entire cell line population (i.e. *n* = 1) or one of multiple clones within a cell line (i.e. *n* ≥ 3). *K*_*i*_, *r*_*i*_ are the carrying capacity and maximum growth rate of each population respectively. *v* is the loss of contact inhibition, assumed to be identical for all clones within a cell line. Two instances of equation (3) were fitted to the data: one representing Richards growth model where *v* was kept variable; the other representing the logistic growth model with *v* := 1.

In addition to the two models mentioned above, we also fitted cell line population growth with the Gompertz model (data not shown). These models were chosen due to their prevalence in the literature describing the growth of tumors and biological interpretation. The growth data was read into R and fit using the package growthrates [45]. In order to compare the quality of growth-model fits we calculate the adjusted-*R*^2^ for each of the three models across all eight cell lines (Supplementary Table 1) We then compared Akaike Information Criterion scores (Supplementary Table 5) and found the Richards model to have the lowest score in 7/8 cell lines. However, identifiability analysis revealed we were unable to confidently infer the growth rate (r) and/or loss of contact inhibition (v) parameters in 4/8 cell lines (Supplementary Fig. 2, Supplementary Table 6). Thus we modeled *in-vitro* growth using the logistic growth model. To model clonal growth dynamics, we used the ODE45 solver for MatLab version 9.7.0.1216025 to solve eq (3) after being parameterized using the values in Supplementary Table 4.

### 4.5 Identifying r/K trade-offs between coexisting clones within a cell line

Let *C* = {*N*_1_, …*N*_*n*_} be the set of *n* clones identified within a given cell line. For each pair {*x, y*|*x, y* ∈ *C* ˄ *x ≠ y*} we use a Student’s t-test to compare the growth rates predicted for cells assigned to clone *x* (*r*_*x*_) vs. cells assigned to clone *y* (*r*_*y*_). We do the same for the carrying capacity predicted for cells of the two clones (*K*_*x*_, *K*_*y*_).

For clone pairs with a *p* - *value* ≤ 0.1 for both *r* and *K* we further calculate 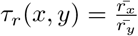 and 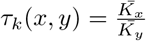. We define (x*,y*) as clones with potential r/K trade-offs:

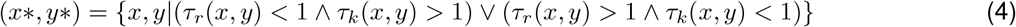

## Notes

### Competing Interest Statement

The authors have declared no competing interest.

### Summary of Updates

Uploaded version of manuscript with old title. Everything else is the same. This one has the correct title: Mathematical modeling of clonal interference by density-dependent selection in heterogeneous cancer cell lines.

