## Supplementary for "Mathematical modeling of clonal interference by density-dependent selection in heterogeneous cancer cell lines"

### Supplementary Information

#### 1 Identifiability analysis

Identifiability analysis is a group of methods used to determine how well the parameters of a model are estimated [25]. The likelihood function is one method that can be used to identify parameter values that are more likely than others. In other words, it tells us which parameter values could be expected to produce the data to which we are fitting the model. Profiling likelihoods is achieved by constructing a function which minimizes the negative log-likelihood for a fixed value of a parameter of interest, and returns $z$, the signed square-root deviance from the minimum. At the maximum likelihood estimation, $z = 0$ by definition.

Let the vector $\theta$ be the group of parameters used in Eqn. 3 to make predictions. Then the likelihood function can be written as $L(\theta|x) = f_\theta(x)$. Suppose that $\theta$ can be decomposed as $\theta = (\delta, \xi)$ where $\delta$ is the parameter of interest and $\xi$ is a vector containing the nuisance parameters. We can re-write our likelihood function as $L(\delta, \xi|x) = f_{\delta,\xi}(x)$. By profiling, we are concentrating the likelihood function for a subset of parameters by expressing the nuisance parameters as a function of the parameter of interest and replacing the nuisance parameter in the likelihood function. So the profile likelihood is then:

$$L_p(\delta) = sup_\xi L(\delta, \xi, x) \tag{5}$$

Let $\epsilon$ be the error between observations and model fit, as defined above, and let growth rate (r) be our parameter of interest, so that $\delta := r$ and $\xi := (K, v)$. Then, in each direction away from $argmax_\delta(L_p(\delta))$, we fix $\delta$ by $\pm s.d.(\epsilon)$, and adjust $\xi$ such that $z$ is minimized. This process continues until $z$ stops changing for new values of $\delta$ or is equal to the MLE fit error. Then, $z$ approximates a chi-squared distribution on which we can compute confidence intervals. We define parameters with confidence intervals that have $>$ 2-fold difference between the 2.5% and 97.5% quantiles as unidentifiable. This analysis revealed that the growth rate (r) or loss of contact inhibition (v) parameters were unidentifiable in Richards model in 4/8 cases (Supplementary Table 6, Supplementary Figure 2). Thus we excluded Richards model, and selected the logistic model given its better fits and lower AIC scores compared to Gompertz (Supplementary Tables 1, 5).

#### 2 Analytical results optimizing cell passaging for logistic growth model (two clone case).

We identified 2 r/K trade-offs between clone pairs across eight cell lines (Fig. 2A, Supplementary Table 4). This suggests that for these cell lines an optimal time for splitting the cells exists, where the trade-off is balanced. Optimizing the timing for splitting cells should thus stabilize a cell line's clonal composition over multiple passages. A condition for such an optimal time to exists is that growth of the population can never be negative. Here we prove that a heterogeneous cell line with a positive growth rate, consisting of two clones with an r/K trade-off, will never have a negative growth rate.

Let $i \in \{1 \dots n\}$ be a clone within a cell line consisting of $n$ clones. Let each clone have its own specific growth rate, $r_i$, and carrying capacity, $K_i$. Then the change in the number of cell members, $N_i$, of clone $i$ can be modeled as:

$$\frac{dN_i}{dt} = (N_i \cdot r_i) \cdot (1 - \frac{\sum_{j=1}^{n} N_j}{K_i}) \tag{6}$$

which has the following solution:

$$N_i(t) = \frac{N_i \cdot K_i}{\sum_{j=1}^{n} N_j + (K_i - \sum_{j=1}^{n} N_j) \cdot e^{-(r_i \cdot t)}} \tag{7}$$

In order to arrive at the solution above in (9), we start by separating the variables in (8), getting our $dt$ on the LHS, and the $dN_i$ on the RHS.

$$(r_i)(dt) = \frac{dN_i}{N_i \cdot (1 - \frac{\sum_{j=1}^{n} N_j}{K_i})} \tag{8}$$

Next, we expand the RHS of (10) through partial fraction decomposition, where $\Gamma$ and $\Psi$ are unknown constants

$$\frac{1}{N_i \cdot (1 - \frac{\sum_{j=1}^{n} N_j}{K_i})} = \frac{\Gamma}{N_i} + \frac{\Psi}{(1 - \frac{\sum_{j=1}^{n} N_j}{K_i})} \tag{9}$$

Multiply both sides of (11) by the LHS denominator to simplify, giving us

$$1 = \Gamma \cdot [1 - \frac{\sum_{j=1}^{n} N_j}{K_i}] + (\Psi \cdot N_i) \tag{10}$$

And so then solving for $\Psi$ and $\Gamma$ we get

$$\Psi = \frac{1}{N_i} - \frac{\sum_{j=1}^{n} N_j}{N_i} + \frac{\Gamma \cdot \sum_{j=1}^{n} N_j}{N_i \cdot K_i} \Rightarrow \Gamma = 1 \tag{11}$$

Now we can return to integrating

$$\int r_i dt = \int \frac{dN_i}{N_i} + \int \frac{\frac{\frac{1}{N_i} - \frac{\sum_{j=1}^{n} N_j}{N_i} + \frac{\Gamma \cdot \sum_{j=1}^{n} N_j}{N_i \cdot K_i}}{K_i}}{1 - \frac{\sum_{j=1}^{n} N_j}{K_i}} \tag{12}$$

The LHS, and first term of RHS are easy enough. For the second term of the RHS, we need to integrate through u-substitution where $u = 1 - \frac{\sum_{j=1}^{n} N_j}{K_i}$ and $du = \frac{-1}{K_i}$ we get the following

$$\int \frac{-du}{u} = -ln[u] = -ln[1 - \frac{\sum_{j=1}^{n} N_j}{K_i}] \tag{13}$$

Then, putting together all terms, and accounting for the constant of integration $C$ we get

$$(r_i \cdot t) + C = \ln[\frac{N_i}{1 - \frac{\sum_{j=1}^{n} N_j}{K_i}}] \tag{14}$$

Next, we exponentiate both sides

$$C \cdot e^{(r_i \cdot t)} = \frac{N_i}{1 - \frac{\sum_{j=1}^{n} N_j}{K_i}} \tag{15}$$

Find how $C = e^C$ relates to initial conditions

$$C = \frac{N_i \cdot K_i}{K_i - \sum_{j=1}^{n} N_j} \tag{16}$$

Plug C into (17) and solve for N

$$N_i(t) = \frac{N_i \cdot K_i}{\sum_{j=1}^{n} N_j + (K_i - \sum_{j=1}^{n} N_j) \cdot e^{-(r_i \cdot t)}} \tag{17}$$

Now, we can move from population sizes to frequencies by letting $p = \frac{N_i}{\sum_{j=1}^{n}(N_j)}$ and $T = \sum_{i=1}^{n}(N_i)$ where $p$ is the proportion of one of our subclones $N_i$, and $T$ is the total population. Thus the equations that describe how $p$ and $T$ change over time, in the two clone case, are as follows:

$$\frac{dp}{dt} = p \cdot (1-p)[r_a \cdot (1 - \frac{T}{K_a}) - r_b \cdot (1 - \frac{T}{K_b})] \tag{18}$$

$$\frac{dT}{dt} = T * [(p * r_a)(1 - \frac{T}{K_a}) + (1-p)(r_b)(1 - \frac{T}{K_b})] \tag{19}$$

where $K_a$ and $K_b$ are the carrying capacity of clone (a) and (b) respectively, and $r_a$ and $r_b$ are the respective growth rates.

Next we sought to prove that equation (21) is always positive. To do so we started out assuming the opposite, that there does exist some time point where $T' < 0$ and arrived at the proof by contradiction. Please note we are most interested in the situation where there is an r/K trade-off. That is, for the rest of this exercise please assume $r_a > r_b$ and $K_a < K_b$. Let's also assume our two populations are well-mixed and initially comprise no more than 60 percent of $max(K_a, K_b)$.

If we let

$$\frac{dT}{dt} = T * [(p * r_a)(1 - \frac{T}{K_a}) + (1-p)(r_b)(1 - \frac{T}{K_b})] = 0 \tag{20}$$

And solve for $T$ we get

$$T = \frac{(K_a * K_b)[(r_b * p) - r_b - (r_a * p)]}{(K_a * r_b * p) - (K_a * r_b) - (K_b * r_a * p)} \tag{21}$$

Equation (23) is the T nullcline. It's easy to see that this function, $T(p)$, is one-to-one. If $T(p_1) = T(p_2)$, then $p_1 = p_2$. The nullcline being a one-to-one function implies that every input which can bring the system to zero, will only bring the system to zero. Now we have $T$ totally in terms of $p$, $r_a$,$r_b$,$K_a$, and $K_b$. Next we can plug it into equation (20) to get it defined in terms of $p$ and the aforementioned $r$ and $K$ parameters:

$$\begin{aligned} \frac{dp}{dt} = \{p * (1 - p)[r_a * (1 - \frac{K_b[(r_b * p) - r_b - (r_a * p)]}{(K_a * r_b * p) - (K_a * r_b) - (K_b * r_a * p)}) - \\ r_b * (1 - \frac{K_a[(r_b * p) - r_b - (r_a * p)]}{(K_a * r_b * p) - (K_a * r_b) - (K_b * r_a * p)})\}] \end{aligned} \tag{22}$$

which is always less than or equal to zero. Thus the system can never cross the nullcline given by $T' = 0$. We know that our system exists in the $T' > 0$ space due to the initial conditions and positive $r$ and $K$ terms, but it can never cross into $T' < 0$ space. This contradicts our original assumption that there existed a timepoint $t$ such that $T'(t) < 0$. Hence $T'$ is always positive. The vector field given by $T'$ and $p'$ with the nullcine $T' = 0$ (Supplementary Fig. 1) shows that the $T$ nullcline passes the Horizontal Line Test, further demonstrating its injective nature. For visualization purposes, the Total Population, $T$ (the x-axis), has been scaled to $max(K_a, K_b) = K_b = 1$. Accordingly, each of our variables, $p$ and $T$, are scaled by $p = p * K_b$ and $T = T * K_b$.

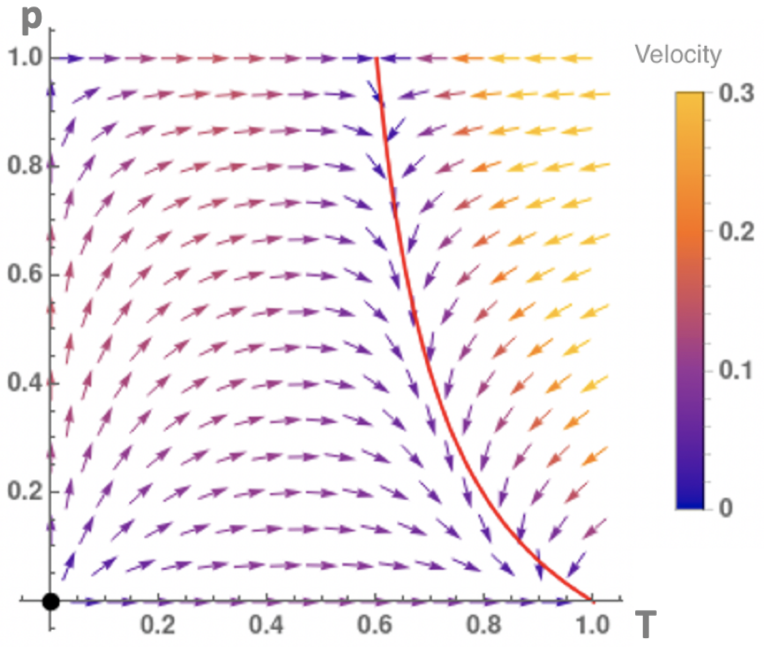


**Supplementary Figure 1: Growth dynamics for the two-clones case**. Vector field given by eqs 20, 21. The red curve shows where there is no change in total population (given by the nullcline $T' = 0$). Arrows indicate total population growth ($T'$). Velocities calculated as $\sqrt{T'(t)^2 + A'(t)^2}$. Velocity is always zero at the nullcline showing that the system can never cross from $T' > 0$ into $T' < 0$ space.

**Supplementary Table 1:** Adjusted-$R^2$ for Model Fits Across Cell Lines.

| | HGC-27 | KATOIII | MKN-45 | NCI-N87 | NUGC-4 | SNU-601 | SNU-638 | SNU-668 |
|---|---|---|---|---|---|---|---|---|
| Logistic | 1.00 | 0.97 | 0.99 | 0.99 | 0.98 | 0.99 | 0.97 | 0.97 |
| Richards | 1.00 | 0.97 | 0.99 | 0.99 | 0.97 | 0.99 | 0.97 | 0.98 |
| Gompertz | 1.00 | 0.95 | 0.96 | 0.95 | 0.98 | 0.98 | 0.95 | 0.98 |

**Supplementary Table 2:** Inferred parameters for best logistic models across cell lines.

| | r | K | bestFit | adjutedRSquared | doublingTime | cellLine |
|---|---|---|---|---|---|---|
| HGC-27 | 0.96 | 5095556.00 | Logistic | 1.00 | 1.10 | HGC-27 |
| KATOIII | 0.44 | 4904316.00 | Logistic | 0.97 | 4.70 | KATOIII |
| MKN-45 | 0.63 | 9237232.00 | Logistic | 0.99 | 3.00 | MKN-45 |
| NCI-N87 | 0.45 | 38306195.00 | Logistic | 0.99 | 2.50 | NCI-N87 |
| NUGC-4 | 0.92 | 6899073.00 | Logistic | 0.98 | 1.10 | NUGC-4 |
| SNU-601 | 0.51 | 7374909.00 | Logistic | 0.99 | 2.30 | SNU-601 |
| SNU-638 | 0.71 | 4791667.00 | Logistic | 0.97 | 1.20 | SNU-638 |
| SNU-668 | 0.45 | 5048178.00 | Logistic | 0.98 | 1.90 | SNU-668 |

**Supplementary Table 3: KEGG pathway biomarkers of growth rate and carrying capacity,.** Adjusted-$R^2$, p-value, and pearson's r for linear models correlating biomarkers with growth parameters. Top five pathways predictive for growth rate and top five pathways predictive for carrying capacity (**bold**) were prioritized during training and further used for validation

| | K: adj.r.squared | K: p.value | K: pearson | r: adj.r.squared | r: p.value | r: pearson |
|---|---|---|---|---|---|---|
| **PI3K-Akt signaling pathway: G0G1** | **0.91** | **0.01** | **-0.97** | -0.21 | 0.61 | 0.31 |
| **PI3K-Akt signaling pathway: all** | **0.72** | **0.04** | **-0.91** | -0.28 | 0.74 | 0.21 |
| **Arginine and proline metabolism: G2M** | **0.62** | **0.07** | **-0.82** | -0.29 | 0.76 | -0.19 |
| **Arginine and proline metabolism: S** | **0.62** | **0.07** | **-0.82** | -0.27 | 0.72 | -0.22 |
| **Amoebiasis: G2M** | -0.33 | 0.94 | 0.11 | **0.89** | **0.01** | **-0.96** |
| **Epstein-Barr virus infection: G0G1** | -0.32 | 0.88 | 0.14 | **0.86** | **0.01** | **-0.95** |
| **Epstein-Barr virus infection: all** | -0.33 | 0.91 | -0.01 | **0.80** | **0.03** | **-0.92** |
| **AMPK signaling pathway: S** | 0.28 | 0.21 | -0.72 | **0.72** | **0.04** | **0.89** |
| AMPK signaling pathway: all | 0.16 | 0.27 | -0.65 | 0.68 | 0.05 | 0.87 |
| AMPK signaling pathway: G0G1 | 0.07 | 0.33 | -0.59 | 0.70 | 0.05 | 0.88 |
| Epstein-Barr virus infection: G2M | -0.29 | 0.78 | -0.10 | 0.66 | 0.06 | -0.87 |
| RNA degradation: G2M | -0.33 | 0.98 | -0.02 | 0.61 | 0.07 | 0.84 |
| Arginine and proline metabolism: all | 0.60 | 0.08 | -0.81 | -0.27 | 0.72 | -0.22 |
| AMPK signaling pathway: G2M | 0.59 | 0.08 | -0.85 | 0.39 | 0.16 | 0.73 |
| Arginine and proline metabolism: G0G1 | 0.55 | 0.09 | -0.79 | -0.25 | 0.68 | -0.25 |
| PI3K-Akt signaling pathway: S | 0.52 | 0.10 | -0.85 | -0.23 | 0.65 | 0.28 |
| Spliceosome: G0G1 | 0.31 | 0.19 | 0.70 | -0.26 | 0.71 | 0.23 |
| Spliceosome: S | 0.31 | 0.19 | 0.69 | -0.27 | 0.72 | 0.22 |
| PI3K-Akt signaling pathway: G2M | 0.31 | 0.19 | -0.72 | -0.33 | 0.98 | -0.02 |
| Pathways in cancer: S | 0.25 | 0.22 | 0.63 | 0.10 | 0.32 | -0.57 |
| Base excision repair: G2M | 0.22 | 0.24 | -0.66 | -0.30 | 0.79 | -0.17 |
| Axon guidance: all | 0.21 | 0.25 | 0.61 | 0.05 | 0.35 | -0.54 |
| Axon guidance: G0G1 | 0.21 | 0.25 | 0.60 | -0.07 | 0.45 | -0.45 |
| Pathways in cancer: G2M | 0.20 | 0.25 | 0.62 | 0.24 | 0.23 | -0.66 |
| Base excision repair: all | 0.20 | 0.25 | -0.65 | -0.25 | 0.69 | -0.25 |
| Spliceosome: all | 0.18 | 0.26 | 0.61 | -0.13 | 0.51 | 0.39 |
| Axon guidance: G2M | 0.08 | 0.33 | 0.54 | 0.15 | 0.29 | -0.60 |
| Base excision repair: G0G1 | 0.07 | 0.34 | -0.54 | -0.02 | 0.41 | -0.49 |
| Pathways in cancer: all | 0.05 | 0.35 | 0.52 | 0.22 | 0.24 | -0.65 |
| Axon guidance: S | 0.03 | 0.36 | 0.52 | 0.42 | 0.14 | -0.75 |
| Pathways in cancer: G0G1 | -0.02 | 0.41 | 0.47 | 0.30 | 0.20 | -0.69 |
| Purine metabolism: all | -0.04 | 0.43 | -0.47 | -0.02 | 0.41 | -0.48 |
| Influenza A: S | -0.05 | 0.44 | 0.41 | -0.05 | 0.44 | -0.46 |
| Proteasome: G0G1 | -0.06 | 0.45 | 0.46 | -0.04 | 0.42 | -0.47 |
| Spliceosome: G2M | -0.07 | 0.46 | 0.43 | -0.07 | 0.45 | 0.44 |
| Purine metabolism: G0G1 | -0.07 | 0.46 | -0.43 | 0.06 | 0.35 | -0.54 |
| Complement and coagulation cascades: S | -0.08 | 0.47 | -0.46 | -0.33 | 0.95 | 0.04 |
| Influenza A: G2M | -0.09 | 0.47 | 0.38 | -0.07 | 0.46 | -0.44 |
| Influenza A: all | -0.09 | 0.47 | 0.38 | -0.02 | 0.40 | -0.49 |
| Purine metabolism: G2M | -0.10 | 0.48 | -0.43 | -0.05 | 0.44 | -0.46 |
| p53 signaling pathway: G2M | -0.10 | 0.48 | 0.36 | 0.26 | 0.22 | 0.67 |
| Pyrimidine metabolism: G2M | -0.10 | 0.48 | -0.41 | 0.08 | 0.33 | -0.56 |
| Influenza A: G0G1 | -0.10 | 0.48 | 0.38 | 0.03 | 0.37 | -0.52 |
| Pyrimidine metabolism: all | -0.14 | 0.53 | -0.38 | 0.08 | 0.33 | -0.56 |
| p53 signaling pathway: S | -0.14 | 0.53 | 0.29 | -0.29 | 0.78 | 0.17 |
| Cell cycle: S | -0.15 | 0.54 | 0.41 | 0.47 | 0.12 | -0.78 |
| Complement and coagulation cascades: G2M | -0.15 | 0.54 | -0.40 | -0.33 | 0.92 | -0.06 |
| Pyrimidine metabolism: G0G1 | -0.15 | 0.54 | -0.35 | 0.19 | 0.26 | -0.63 |
| Purine metabolism: S | -0.16 | 0.55 | -0.37 | 0.00 | 0.39 | -0.50 |

| | | | | | | |
|---|---|---|---|---|---|---|
| Complement and coagulation cascades: all | -0.16 | 0.56 | -0.38 | -0.32 | 0.90 | -0.08 |
| Hepatitis B: G2M | -0.17 | 0.56 | 0.31 | 0.01 | 0.38 | -0.51 |
| Epstein-Barr virus infection: S | -0.17 | 0.56 | -0.29 | 0.45 | 0.13 | -0.77 |
| Complement and coagulation cascades: G0G1 | -0.18 | 0.58 | -0.35 | -0.32 | 0.86 | -0.11 |
| p53 signaling pathway: all | -0.19 | 0.59 | 0.24 | -0.21 | 0.62 | 0.30 |
| p53 signaling pathway:G0G1 | -0.20 | 0.61 | 0.23 | -0.22 | 0.63 | 0.30 |
| Hippo signaling pathway: G2M | -0.21 | 0.62 | 0.32 | 0.12 | 0.30 | -0.59 |
| Hepatitis B: S | -0.21 | 0.62 | 0.26 | -0.05 | 0.43 | -0.46 |
| Cell cycle: all | -0.21 | 0.62 | 0.39 | 0.31 | 0.19 | -0.70 |
| Pertussis: all | -0.21 | 0.63 | -0.31 | 0.08 | 0.33 | -0.55 |
| Hippo signaling pathway: G0G1 | -0.22 | 0.63 | 0.31 | 0.15 | 0.28 | -0.60 |
| Pertussis: G0G1 | -0.22 | 0.63 | -0.30 | 0.08 | 0.33 | -0.55 |
| Hippo signaling pathway: S | -0.23 | 0.64 | 0.28 | 0.09 | 0.32 | -0.57 |
| Small cell lung cancer: S | -0.23 | 0.64 | 0.21 | -0.33 | 0.93 | -0.06 |
| Proteasome: S | -0.23 | 0.64 | -0.20 | 0.22 | 0.24 | -0.64 |
| Hippo signaling pathway: all | -0.23 | 0.64 | 0.29 | 0.10 | 0.32 | -0.57 |
| Ribosome biogenesis in eukaryotesG2M | -0.23 | 0.65 | -0.23 | -0.24 | 0.67 | 0.26 |
| Pertussis: S | -0.24 | 0.66 | -0.28 | 0.11 | 0.31 | -0.57 |
| Proteasome: all | -0.24 | 0.66 | 0.31 | 0.00 | 0.39 | -0.50 |
| Small cell lung cancer: G2M | -0.26 | 0.70 | 0.18 | -0.28 | 0.74 | -0.20 |
| Pyrimidine metabolism: S | -0.26 | 0.71 | -0.23 | 0.20 | 0.25 | -0.63 |
| ECM-receptor interaction: G0G1 | -0.27 | 0.72 | -0.31 | -0.03 | 0.42 | 0.48 |
| ECM-receptor interaction: S | -0.27 | 0.73 | -0.30 | -0.08 | 0.47 | 0.43 |
| RNA transport: S | -0.28 | 0.74 | 0.17 | 0.46 | 0.13 | 0.77 |
| Hepatitis B: all | -0.28 | 0.75 | 0.15 | -0.04 | 0.42 | -0.47 |
| Pertussis: G2M | -0.28 | 0.76 | -0.20 | 0.15 | 0.29 | -0.60 |
| Amyotrophic lateral sclerosis (ALS): G0G1 | -0.29 | 0.76 | -0.18 | -0.21 | 0.62 | -0.31 |
| ECM-receptor interaction: all | -0.29 | 0.78 | -0.26 | -0.13 | 0.51 | 0.39 |
| Hepatitis B: G0G1 | -0.30 | 0.80 | 0.12 | -0.01 | 0.40 | -0.49 |
| Base excision repair: S | -0.30 | 0.81 | -0.18 | -0.17 | 0.57 | -0.35 |
| RNA transport: G0G1 | -0.30 | 0.81 | 0.12 | 0.51 | 0.11 | 0.79 |
| Amyotrophic lateral sclerosis (ALS): all | -0.30 | 0.81 | -0.15 | -0.18 | 0.58 | -0.34 |
| RNA transport: all | -0.31 | 0.84 | 0.09 | 0.56 | 0.09 | 0.82 |
| Amoebiasis: G0G1 | -0.32 | 0.85 | -0.03 | 0.25 | 0.22 | -0.66 |
| RNA degradation: G0G1 | -0.32 | 0.87 | 0.07 | 0.48 | 0.12 | 0.78 |
| Small cell lung cancer: all | -0.32 | 0.89 | 0.03 | -0.33 | 0.91 | -0.07 |
| Cell cycle: G2M | -0.32 | 0.89 | 0.14 | -0.31 | 0.82 | 0.15 |
| Ribosome biogenesis in eukaryotes: S | -0.32 | 0.89 | -0.03 | -0.33 | 0.91 | -0.07 |
| Amyotrophic lateral sclerosis (ALS): G2M | -0.32 | 0.89 | -0.08 | -0.13 | 0.51 | -0.39 |
| RNA degradation: all | -0.32 | 0.90 | 0.05 | 0.52 | 0.10 | 0.80 |
| Ribosome biogenesis in eukaryotes: all | -0.33 | 0.91 | -0.02 | -0.33 | 0.93 | 0.06 |
| Amoebiasis: S | -0.33 | 0.91 | 0.01 | 0.56 | 0.09 | -0.82 |
| Cell cycle: G0G1 | -0.33 | 0.92 | 0.15 | 0.25 | 0.22 | -0.66 |
| Amyotrophic lateral sclerosis (ALS): S | -0.33 | 0.92 | -0.06 | -0.12 | 0.50 | -0.40 |
| RNA transport: G2M | -0.33 | 0.93 | 0.03 | 0.52 | 0.10 | 0.80 |
| Amoebiasis: all | -0.33 | 0.93 | 0.03 | 0.55 | 0.09 | -0.81 |
| ECM-receptor interaction: G2M | -0.33 | 0.94 | -0.14 | -0.26 | 0.70 | 0.24 |
| Small cell lung cancer: G0G1 | -0.33 | 0.96 | -0.01 | -0.29 | 0.76 | -0.19 |
| Ribosome biogenesis in eukaryotes: G0G1 | -0.33 | 0.99 | 0.05 | -0.33 | 0.97 | -0.03 |
| Proteasome: G2M | -0.33 | 0.99 | 0.05 | -0.03 | 0.42 | -0.48 |
| RNA degradation: S | -0.33 | 0.99 | -0.02 | 0.60 | 0.08 | 0.84 |

| | K | r | CL |
|---|---|---|---|
| 102944 | 4690105 | 0.49 | SNU-668 |
| 102945 | 4674250 | 0.38 | SNU-668 |
| 102948 | 3450777 | 0.53 | SNU-668 |
| 102950 | 4331258 | 0.66 | SNU-668 |
| 102951 | 5488043 | 0.53 | SNU-668 |
| 102952 | 4379298 | NA | SNU-668 |
| 102954 | 5046408 | 0.62 | SNU-668 |
| 102955 | 4773230 | 0.52 | SNU-668 |
| 106394 | 3166668 | 0.56 | KATOIII |
| 106396 | 3117464 | NA | KATOIII |
| 106399 | 3274342 | 0.53 | KATOIII |
| 106404 | 3346502 | 0.58 | KATOIII |
| 112380 | 7448893 | 0.32 | SNU-601 |
| 112382 | 8148176 | 0.34 | SNU-601 |
| 112387 | 7231439 | NA | SNU-601 |
| 112389 | 6627640 | 0.32 | SNU-601 |
| 112392 | 6798104 | 0.34 | SNU-601 |
| 112399 | 6312316 | NA | SNU-601 |
| 112402 | 8050488 | 0.38 | SNU-601 |
| 112404 | 7081546 | 0.31 | SNU-601 |
| 112408 | 8255791 | 0.37 | SNU-601 |
| 112410 | 7903615 | NA | SNU-601 |
| 112413 | 7196330 | 0.32 | SNU-601 |
| 114525 | 11055814 | 0.65 | MKN-45 |
| 114530 | 10785452 | 0.68 | MKN-45 |
| 119963 | 18156049 | NA | NCI-N87 |
| 119965 | 22002556 | 0.62 | NCI-N87 |
| 119967 | 20354070 | 0.57 | NCI-N87 |
| 119968 | 18548305 | 0.67 | NCI-N87 |
| 122360 | 5156522 | 0.80 | SNU-638 |
| 122361 | 4331370 | NA | SNU-638 |
| 122363 | 4759109 | 0.80 | SNU-638 |
| 125616 | 9123026 | 0.78 | NUGC-4 |
| 125618 | 9655395 | 0.69 | NUGC-4 |
| 125619 | 8845460 | 0.82 | NUGC-4 |
| 129343 | 9429235 | 0.85 | HGC-27 |
| 129344 | 8996528 | 0.82 | HGC-27 |
| 129345 | 8856699 | 0.76 | HGC-27 |
| 129346 | 8616362 | 0.78 | HGC-27 |

**Supplementary Table 4:** Predicted growth parameters for clones (rows) identified in eight gastric cancer lines. Parameter predictions come from linear models built between pathway biomarkers and inferred parameter values at the cell line level. The input is clonal pathway activity, and the output is the predicted growth parameter for that clonal population.

**Supplementary Table 5: Difference in AIC scores across cell lines**. Difference AIC scores were calculated using the dAIC function from the package bbmle for the statistical programming language R. For 7/8 cell lines, the Richards model has the lowest AIC score. Thus the added parsimony can be justified by the increase in goodness of fit.

| Difference in Akaike Information Criterion scores across cell lines | | | | | | | | |
|---|---|---|---|---|---|---|---|---|
| dAIC | HGC-27 | KATOIII | MKN-45 | NCI-N87 | NUGC-4 | SNU-601 | SNU-638 | SNU-668 |
| Richards (3 params) | 0 | 0 | 0 | 0 | 1.0 | 0 | 0 | 0 |
| Logistic (2 params) | 151.4 | 54.6 | 22.1 | 17 | 7.8 | 3.8 | 16.8 | 77 |
| Gompertz (2 params) | 86.5 | 116.8 | 101.2 | 34.2 | 0 | 105.6 | 36.6 | 1.3 |

**Supplementary Table 6: Likelihood profile confidence intervals across paramters and model types**. Confidence intervals generated from likelihood profiles for all parameters in Richards and logistic (Verhulst) growth models. Entries with >2 fold difference between the 2.5% and 97.5% quantiles highlighted in red. Growth rate (r) is not identifiable for NUGC-4 or SNU-668 in the Richards model. Additionally, loss of contact inhibition (v) is not identifiable for those two cell lines, NCI-N87, and SNU-638.

| Likelihood profile confidence intervals | | | | | |
|---|---|---|---|---|---|
| | | **Richards** | | **Logistic** | |
| **Cell Line** | **Parameter** | 2.50% | 97.50% | 2.50% | 97.50% |
| HGC-27 | K | 0.69 | 0.70 | 0.66 | 0.67 |
| KATOIII | K | 0.64 | 0.66 | 0.66 | 0.69 |
| MKN-45 | K | 1.19 | 1.24 | 1.25 | 1.30 |
| NCI-N87 | K | 4.90 | 5.15 | 5.07 | 5.51 |
| NUGC-4 | K | 0.92 | 0.96 | 0.91 | 0.95 |
| SNU-601 | K | 0.96 | 1.00 | 0.99 | 1.02 |
| SNU-638 | K | 0.59 | 0.64 | 0.64 | 0.71 |
| SNU-668 | K | 0.69 | 0.73 | 0.64 | 0.67 |
| HGC-27 | r | 1.77 | 2.31 | 1.04 | 1.08 |
| KATOIII | r | 0.24 | 0.26 | 0.33 | 0.36 |
| MKN-45 | r | 0.45 | 0.49 | 0.53 | 0.55 |
| NCI-N87 | r | 0.30 | 0.33 | 0.37 | 0.41 |
| NUGC-4 | r | **1.43** | **75.46** | 0.97 | 1.06 |
| SNU-601 | r | 0.43 | 0.47 | 0.47 | 0.48 |
| SNU-638 | r | 0.37 | 0.43 | 0.52 | 0.59 |
| SNU-668 | r | **1.41** | **158.72** | 0.55 | 0.59 |
| HGC-27 | v | 0.30 | 0.43 | - | - |
| KATOIII | v | 2.14 | 3.17 | - | - |
| MKN-45 | v | 1.39 | 1.98 | - | - |
| NCI-N87 | v | **1.81** | **5.78** | - | - |
| NUGC-4 | v | **0.01** | **0.64** | - | - |
| SNU-601 | v | 1.04 | 1.48 | - | - |
| SNU-638 | v | **2.14** | **5.80** | - | - |
| SNU-668 | v | **0.02** | **0.23** | - | - |

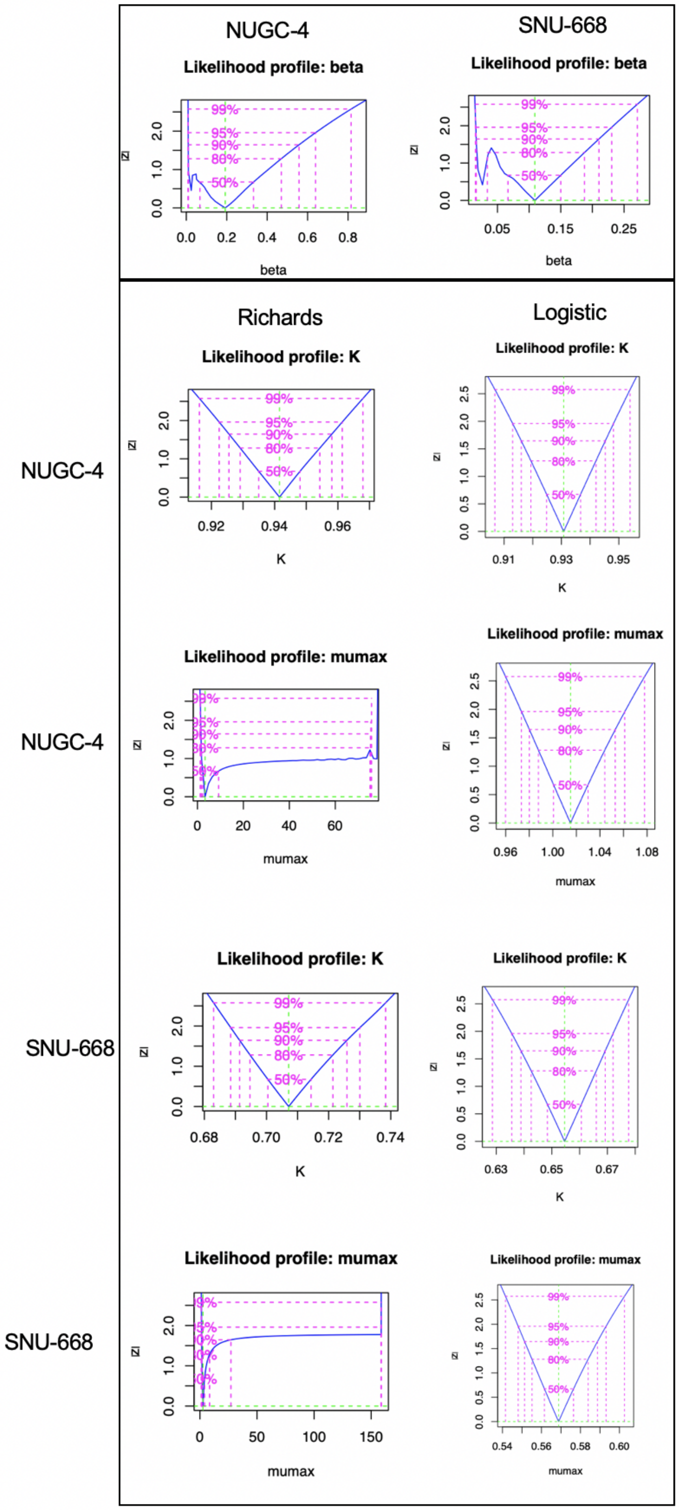


**Supplementary Figure 2: Likelihood profiles across cell lines.** Graphical representation of the likelihood profiles for both parameters in the logistic growth model for all cell lines. Please note here mumax is the growth rate (r). Maximum likelihood values are at the center of the x-axis, with the y-axis giving the absolute value of the z-statistic from the chi-square distribution these profiles approximate. Confidence intervals given at various percentages. Tight confidence intervals tell us when parameters are identifiable.

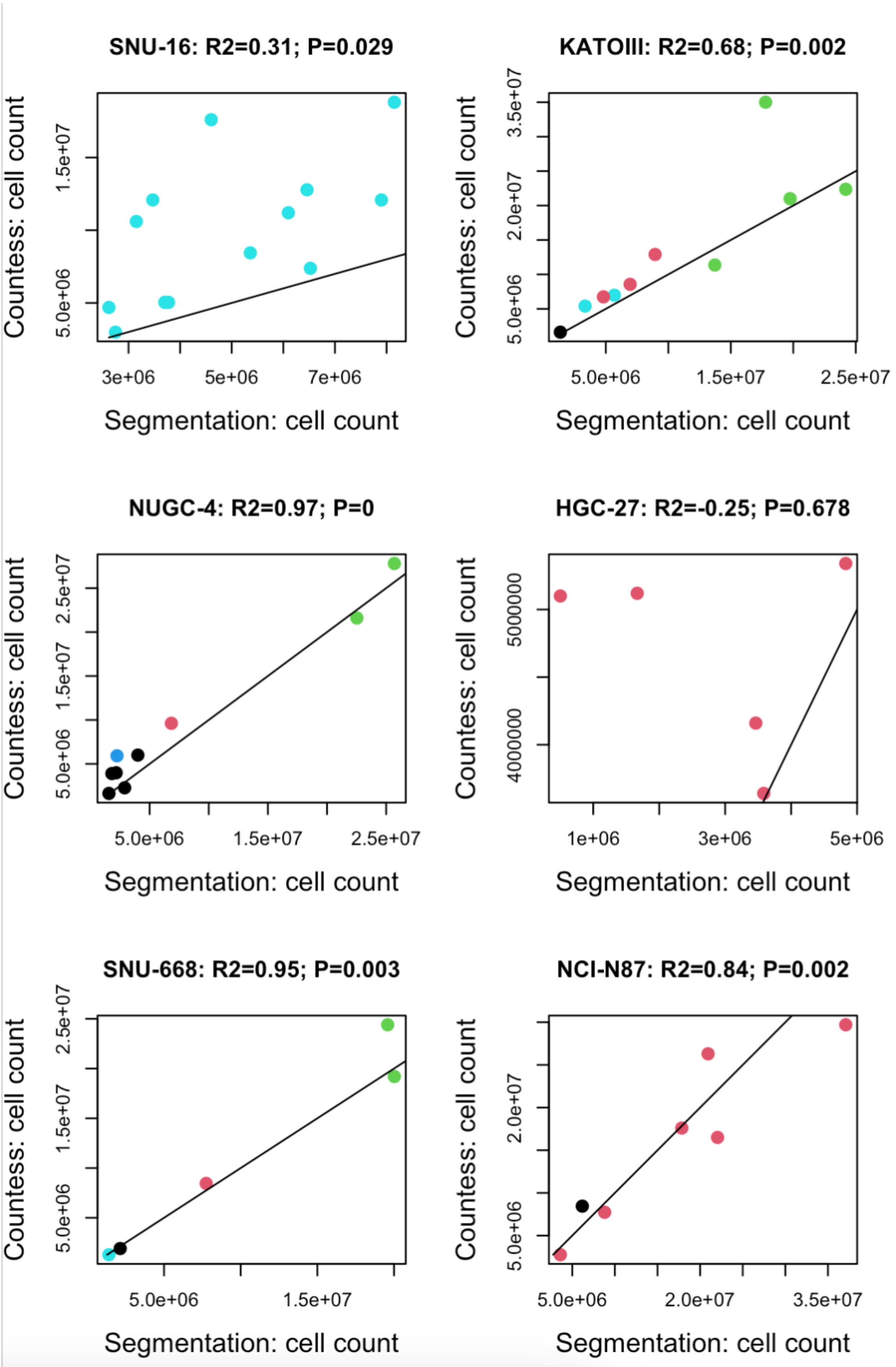


**Supplementary Figure 3: Segmentation evaluation using Countess.** Comparing Countess derived cell counts to cell counts derived from live cell imaging for each cell line.
